# A simple Turing reaction-diffusion model can explain how mother centrioles break symmetry to generate a single daughter

**DOI:** 10.1101/2023.02.02.526828

**Authors:** Zachary M. Wilmott, Alain Goriely, Jordan W. Raff

## Abstract

Centrioles duplicate when a mother centriole gives birth to a daughter that grows from its side. Polo-like-kinase 4 (PLK4), the master regulator of centriole duplication, is recruited symmetrically around the mother centriole, but it then concentrates at a single focus that defines the daughter centriole assembly site. How PLK4 breaks symmetry is unclear. Here, we propose that phosphorylated and unphosphorylated species of PLK4 form the two components of a classical Turing reaction-diffusion system. These two components bind-to/unbind-from the surface of the mother centriole at different rates, allowing a slow-diffusing activator species of PLK4 to accumulate at a single site on the mother, while a fast-diffusing inhibitor species of PLK4 suppresses activator accumulation around the rest of the centriole. This “short-range activation/long-range inhibition”, inherent to Turing-systems, can drive PLK4 symmetry breaking on a continuous centriole surface, with PLK4 overexpression producing multiple PLK4 foci and PLK4 kinase inhibition leading to uniform PLK4 accumulation—as observed experimentally.

## Introduction

Most human cells are born with a single pair of centrioles comprising an older ‘mother’ and a younger ‘daughter’ (Figure 1A); these organelles play an important part in many aspects of cellular organisation (Nigg and Raff, 2009; Bettencourt-Dias et al., 2011; Bornens, 2021; LeGuennec et al., 2021). The centriole pair duplicates precisely once during each cell division cycle when the original centriole pair separate, and a single new daughter grows off the side of each pre-existing centriole (now both termed mothers) (Fırat-Karalar and Stearns, 2014; Nigg and Holland, 2018; Banterle and Gönczy, 2017). The centriole is a 9-fold symmetric structure and it is unclear how its symmetry is broken to establish the single site for daughter centriole assembly. Polo-like-kinase 4 (PLK4) is the master regulator of centriole biogenesis (Habedanck et al., 2005; Bettencourt-Dias et al., 2005), and it appears to be initially recruited around the entire surface of the mother centriole before it becomes concentrated at a single focus that defines the daughter centriole assembly site (Figure 1B) (Arquint and Nigg, 2016; Nigg and Holland, 2018; Gönczy and Hatzopoulos, 2019; Yamamoto and Kitagawa, 2021).

**Figure 1.**
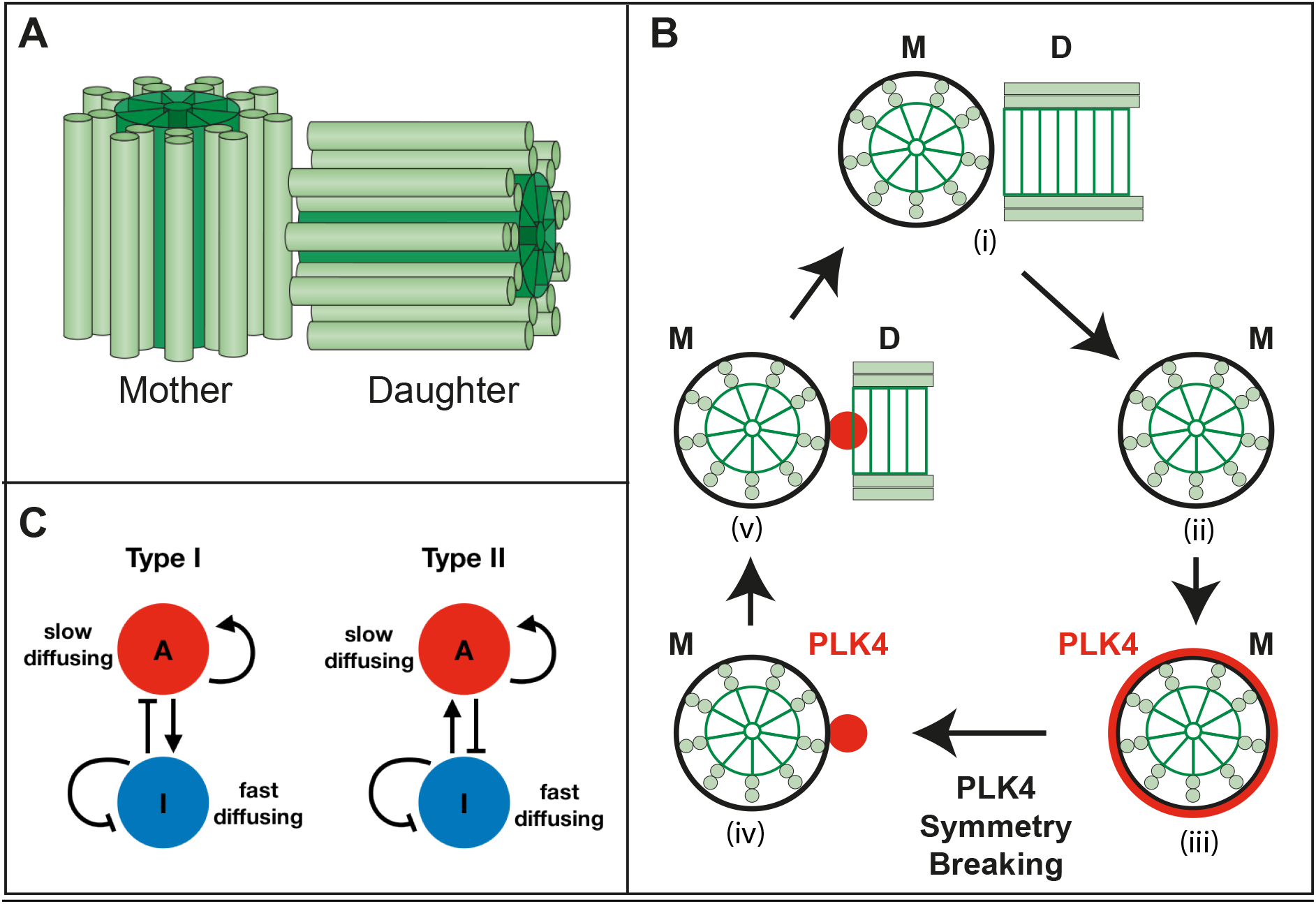
Schematic illustration of centrioles, centriole duplication, PLK4 symmetry breaking, and Turing reaction-diffusion systems. **(A)** Most higher eukaryotic cells are born with a single pair of centrioles comprising an older mother and a younger daughter formed around a 9-fold symmetric central cartwheel (*dark green*) surrounded by 9-sets of microtubule doublets or triplets (*light green*), depending on the species. The centrioles depicted here are from *Drosophila*, which have doublet MTs and which lack the distal and sub-distal appendages found on human mother centrioles. **(B)** The centriole pair (i) duplicates once each cell cycle when the mother (M) and daughter (D) separate (ii) (for simplicity only the original mother is shown here); the daughter matures into a mother (not shown) and both mother centrioles recruit PLK4 (*red*) symmetrically around themselves (iii). PLK4 symmetry is broken (iv) when PLK4 becomes concentrated in a single spot that defines the site of new daughter centriole assembly (v). PLK4 usually dissociates from the centriole by the time daughter centriole assembly is complete (Aydogan et al., 2020). **(C)** Diagrams illustrate the logic of the two chemical reaction schemes that can form Turing reaction-diffusion systems (see text for details).

Two different mathematical models have previously been proposed to explain how PLK4 symmetry might be broken. The approach developed by Kitagawa and colleagues (Takao et al., 2019), was motivated by the observation that PLK4 has an intrinsic ability to self-organise into macromolecular condensates in vitro (Montenegro Gouveia et al., 2018; Yamamoto and Kitagawa, 2019; Park et al., 2019), and that PLK4 and its centriole receptor CEP152 initially appear to be organised into discrete compartments around the centriole—∼12 for CEP152 and ∼6 for PLK4 (Takao et al., 2019). The authors assume that PLK4 initially binds equally well to the twelve CEP152 compartments and that, due to its self-assembling properties, this PLK4 then recruits additional PLK4. They allow PLK4 to autoactivate itself as the local concentration of PLK4 rises, and suppose that active PLK4 is more mobile in the condensates than inactive PLK4 (as observed experimentally). Thus, the active PLK4 generated in one compartment can activate PLK4 in nearby compartments, which stimulates the PLK4 to leave the nearby compartment. In this way, each of the ∼12 CEP152 compartments effectively compete to recruit PLK4, which can then “laterally inhibit” recruitment at nearby compartments. This process allows the ∼12 CEP152 compartments to generate a “pre-pattern” of ∼6 PLK4 compartments. At the start of S-phase, the addition of STIL into the system generates additional positive feedback and ensures that the pre-patterned site with most PLK4 becomes the single dominant site.

In an alternative approach developed by Goryachev and colleagues (Leda et al., 2018), the centriolar surface is also modelled as a number of discrete compartments but, unlike the Takao et al. model, these are not spatially ordered (so all compartments are equivalent and any chemical species produced at one compartment can diffuse equally to all the other compartments). These compartments bind cytoplasmic PLK4, but also the key centriole-assembly protein STIL. PLK4 and STIL interact both in the cytoplasm and in the centriole compartments, with exchange between the centriole compartments occurring via the shared cytoplasm. The phosphorylation of STIL by PLK4 is postulated to increase the centriolar retention of the phosphorylated PLK4:STIL complexes. Since PLK4 can promote its own activation (Lopes et al., 2015; Moyer et al., 2015; Klebba et al., 2015), this system forms a positive feedback loop in which PLK4 activity locally auto-amplifies itself by phosphorylating PLK4:STIL complexes to strengthen their centriolar retention. This process creates a competition between the different compartments for binding to the various PLK4:STIL complexes. This system can break symmetry to establish a single dominant compartment that concentrates the phosphorylated PLK4:STIL complexes.

The different mathematical frameworks used in the two models make it difficult to compare them, so it is unclear if there are any similarities in the logic of the underlying biochemical reactions that determine the behaviour of each system that can explain how two such apparently different models can both break symmetry. Moreover, although both models can account for certain aspects of PLK4 symmetry breaking, neither provide a complete description. For example, the 12-fold symmetry of the CEP152 Receptor and 6-fold symmetry of the PLK4 pre-pattern described by Takao et al., have not been observed in other systems (Tian et al., 2021; Wainman, 2021; Gao et al., 2021; Wilmerding et al., 2023). This model also explicitly relies on STIL appearing in the system only after PLK4 has already broken symmetry; this may be plausible in somatic cells, but not in rapidly dividing systems such as the early *Drosophila* embryo where the cytoplasmic concentration of Ana2/STIL remains constant through multiple rounds of centriole duplication (Steinacker et al., 2022). The Leda et al. model predicts that inhibiting PLK4 kinase activity should deplete PLK4 from the centriole surface, but it is now clear that PLK4 accumulates at centrioles when its kinase activity is inhibited (Ohta et al., 2018; Yamamoto and Kitagawa, 2019).

In an attempt to address these issues, we have modelled PLK4 symmetry breaking as a “Turing system”, which we define here as a two-component reaction-diffusion system that breaks symmetry through activator-inhibitor dynamics (Turing, 1952; Gierer and Meinhardt, 1972). Turing systems are famous for their ability to produce complex spatial patterns, and they have been well-studied in relation to numerous biological phenomena including the formation of animal coat patterns, predator-prey dynamics, and the spread of disease (Bard, 1981; Levin and Segel, 1985; Mimura and Murray, 1978; Sun, 2012). These systems rely on “short-range activation/long-range inhibition”, whereby relatively fast chemical reactions between the two components initially drive the system towards a steady state, which can then be destabilised by the differential diffusion of the two components to drive pattern-forming instabilities over a longer time-scale.

Although the two previous models of PLK4 symmetry breaking appear to be very different, we show that both can be reformulated as Turing systems, with phosphorylated/non-phosphorylated species of PLK4 (either on its own, or in a complex with STIL) forming the two components in the system that bind/unbind from centrioles at different rates (allowing the two components to effectively differentially diffuse within the system). Through computer simulations we demonstrate that these simple models can break PLK4 symmetry to form a single PLK4 peak, while overexpressing PLK4 can lead to the formation of multiple PLK4 peaks, and inhibiting PLK4 kinase activity can lead to the uniform accumulation of PLK4 around the centriole surface—as observed experimentally. Unexpectedly, our analysis reveals that, in the appropriate parameter regime, the dominant chemical reactions that drive symmetry breaking in the two previous models are actually the same.

Potentially importantly, and in contrast to the previous models, our models do not require that discrete PLK4-binding compartments are already present on the centriole surface prior to PLK4 binding and subsequent symmetry breaking. Although our models can work on a centriole surface comprising any number of discrete PLK4-binding compartments, they do not require this, and they can break PLK4 symmetry even on a continuous centriole surface.

## Results

Throughout this paper, we model the recruitment of PLK4 around the surface of the centriole as a two-component reaction-diffusion system acting on a one-dimensional ring (the centriole surface). It may be shown mathematically that the probabilistic random walk of a molecule that repeatedly binds to, and then unbinds from, a surface on the microscale acts as a diffusive process on the macroscale (Skellam, 1973). Therefore, we use surface diffusion as a simplifying modelling approximation for diffusion through the bulk, where the effective diffusion rate of a molecule along the surface (in our case the centriole surface) is directly related to the rate at which the molecule binds-to/unbinds-from that surface. In such a system, a significant proportion of the molecules that unbind from the surface will inevitably diffuse away rather than rebind to the centriole surface. Calculating the “return probability” of diffusing species in such a system is a complex issue (Revelli et al., 2007; Chechkin et al., 2012), however we shall make the simplifying assumption that this “loss” term may be absorbed into the reaction components of the equations (see below).

The two components in such systems are always a slowly diffusing (i.e. rapidly binding and/or slowly unbinding) species termed an Activator (*A*), and a rapidly diffusing (i.e. slowly binding and/or rapidly unbinding) species termed an Inhibitor (*I*), which satisfy

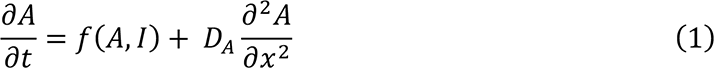

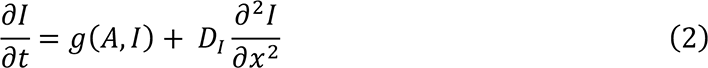

where *t* is time, *x* is arc length (i.e. position) along the circular centriole surface, *f* and *g* are prescribed functions that describe the chemical reactions that determine how *A* and *I* accumulate and decay over time around the centriole surface, and *D_A_* and *D_I_* are the diffusivities of *A* and *I* on the centriole surface, respectively. We use this model as our basic framework.

In order for such a general reaction-diffusion system to break symmetry (i.e. to be a Turing system), there are two conditions that the model must satisfy (Ruan, 1998). First, there must exist a steady state (independent of *x* and *t*) that is stable in the absence of diffusion. Second, this steady state must be unstable in the presence of diffusion. Such a system breaks symmetry as the relatively fast chemical reactions initially drive the system towards the steady state, but this becomes destabilised by diffusion, which drives the pattern-forming instability over a longer time-scale.

These conditions put constraints on the values of the equation parameters that can support symmetry breaking in Turing-systems. Mathematically, these constraints can be expressed for any given system as a set of inequalities that define the possible range of parameter values that will support symmetry breaking. As a consequence of these constraints, symmetry breaking requires that the activator and inhibitor in these systems only interact with each other in one of two well-defined regimes (Type I and Type II, Figure 1C). In both regimes, the inhibitor must diffuse more rapidly than the activator in order to drive a process known as ‘short-range activation, long-range inhibition’. It is possible to generate (non-Turing) two component systems that break symmetry without invoking differential diffusion (Chau et al., 2012). In such cases, symmetry can be broken if one of the two components can be depleted from the cytoplasm. Such cytoplasmic depletion is almost certainly not occurring during PLK4 symmetry breaking — FRAP experiments, for example, show that PLK4 continuously turns over at centrioles (Cizmecioglu et al., 2010; Yamamoto and Kitagawa, 2019). An in-depth derivation and analysis of the stability criteria for Turing-type activator-inhibitor systems can be found in (Ruan, 1998), and we summarise the key results in Appendix I.

### Model 1: Activator to Inhibitor conversion based on the model of Takao et al., 2019

The first model we analyse is a Type I system in which a slowly-diffusing activator *A* is converted into a rapidly-diffusing inhibitor *I* via phosphorylation (Figure 2A). As a biological example, we adapt the model originally proposed by Takao et al. (2018), in which unphosphorylated, kinase inactive, PLK4 is initially recruited to the centriole surface where it self-assembles into slowly turning-over macromolecular condensates. As PLK4 levels in the condensates increase, Takao et al. allow PLK4 to auto-phosphorylate itself to create PLK4*, which turns-over more rapidly within the condensates, as observed in vitro (Montenegro Gouveia et al., 2018; Yamamoto and Kitagawa, 2019; Park et al., 2019). This difference in the turn-over rates of the non-phosphorylated and phosphorylated species is the basis for the differential diffusion of the two components in the Turing-system we formulate here. Takao et al. assume that the non-phosphorylated PLK4 self-assembly rate is subject to a sigmoidal attenuation (i.e. as the condensates grow, they become less likely to disassemble) because the central regions of the condensate become progressively more isolated from the cytoplasm as the condensate grows. We also adopt this sigmoidal self-assembly relationship for the production of *A* (*red box*, equation 3). Whereas Takao et al. assume that PLK4 (*A*) and PLK4* (*I*) are restricted to binding to a defined number of discrete, spatially-organised compartments around the periphery of the centriole, in our model we allow both species to bind freely anywhere on a continuous centriole surface. This model reads:

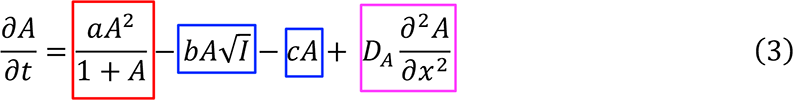

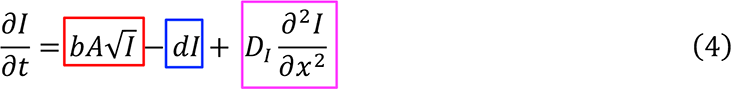

**Figure 2.**
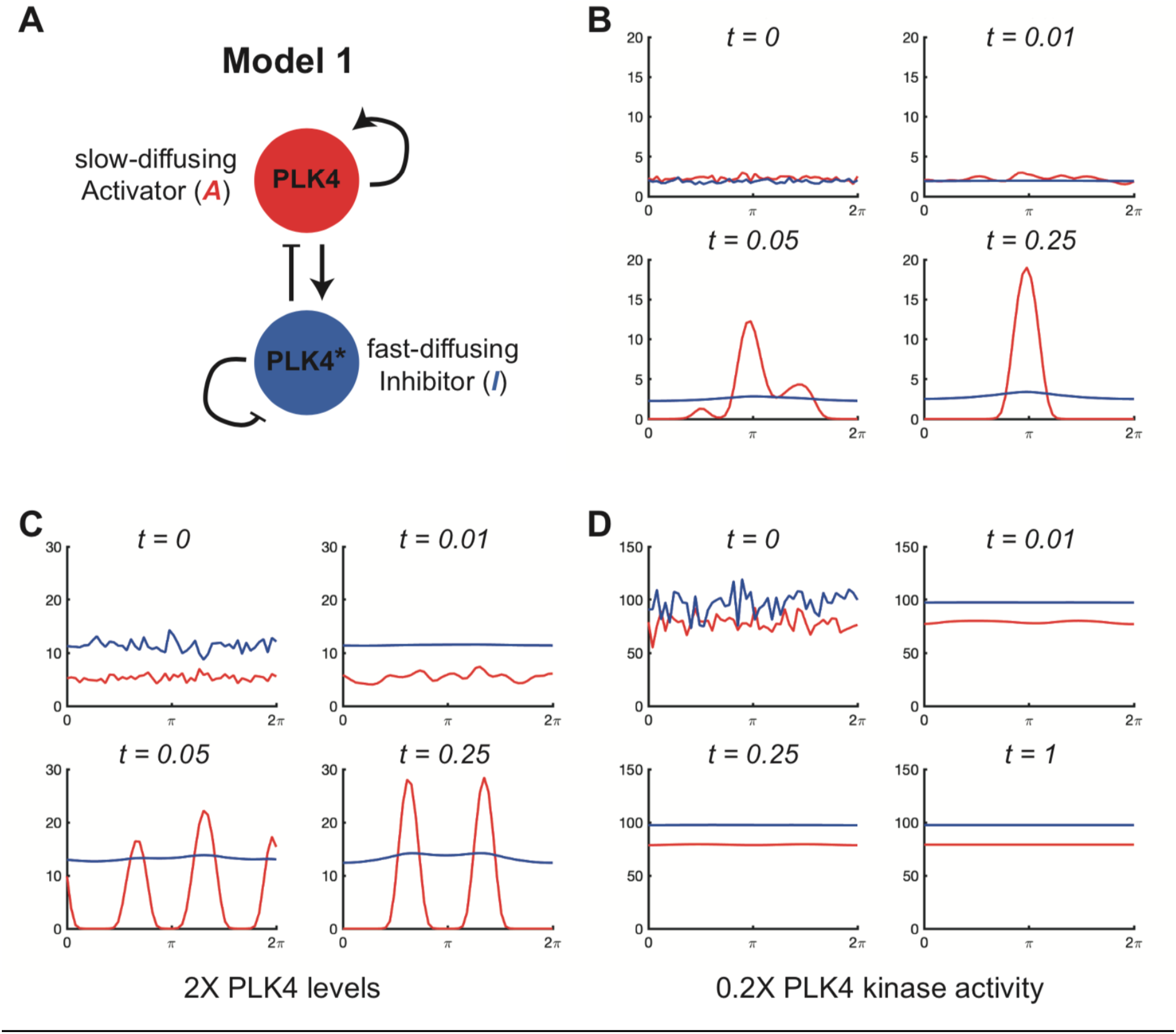
Computer simulations of mathematical Model 1. **(A)** Schematic summarises the reaction regime of Model 1, a Type I Turing-system. The biology underlying this model was originally proposed by Takao et al., (2019). Although it is not possible to precisely assign the activator and inhibitor relationships depicted in this schematic to specific terms in the reaction equations (equations 3 and 4, main text) we can approximately see that *A* (non-phosphorylated PLK4, *red*) promotes the production of more *A* through self-assembly, and the production of *I* (phosphorylated PLK4, denoted PLK4*, *blue*) by stimulating the self-phosphorylation of *A* as the concentration of *A* increases. *I* inhibits the production of *A* by converting it to *I* via phosphorylation, and *I* inhibits its own accumulation by promoting its own degradation/dissociation. **(B-D)** Graphs show how the levels of the activator species (*red* lines) and the inhibitor species (*blue* lines) change over time (arbitrary units) at the centriole surface (modelled as a continuous ring stretched out as a line between 0-2ρχ) in computer simulations. The rate parameters for each simulation are defined in the main text and can lead to symmetry breaking to a single peak (B), or to multiple peaks when cytoplasmic PLK4 levels are increased (C), or to no symmetry breaking at all (with the accumulation of relatively high-levels of activator and inhibitor species spread uniformly around the centriole surface) when PLK4 kinase activity is reduced (D).

The boxed parts of the equation describe how *A* (eq. 3) or *I* (eq. 4) are produced (*red*), removed (*blue*) or diffuse (*magenta*) within the system. The rate parameters *a*, *b*, *c*, and *d* correspond to the self-assembly rate of the unphosphorylated PLK4 complex (i.e. the rate at which *A* is produced in the system) (*a*), the phosphorylation rate of PLK4 by phosphorylated PLK4* (i.e. the rate at which *I* converts *A* into *I*) (*b*), and the rate at which unphosphorylated PLK4 (*c*) or phosphorylated PLK4* (*d*) are either degraded or lost to the cytoplasm. The precise functional form of these equations, which correspond to the strength of the self-assembly of PLK4 and the strength of trans-autophosphorylation of inactive PLK4 by active PLK4*, respectively, are discussed in Appendix II.

In general, it is not possible to precisely assign the activator and inhibitor relationships depicted in the schematic in Figure 2A to specific terms in the reaction equations (eq.3 and eq.4). This is because, mathematically, a positive feedback for *A* means that 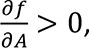 and a negative feedback for *I* means that 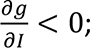 these inequalities may be achieved with complex expressions that extend beyond the usual proportional relationships often assumed. Nevertheless, an approximate description is provided in the Figure legend. As discussed above, the conditions for this system to break symmetry take the form of a set of inequalities for the parameters *a*, *b*, c, and *d*, which we derive in Appendix II. Unless stated otherwise, we choose the parameter values *a* = 500, *b* = 250, *c* = 0, *d* = 400, *D_A_* = 2, and *D_I_* = 5000, that satisfy these conditions. These values were chosen in part to reflect the parameter values and ratios used in the previous modelling papers (Leda et al., 2018; Takao et al., 2019) when adjusted to the same timescale.

In Figure 2B, we plot the solution of this model (i.e. a computer simulation of how PLK4 (*A*) and PLK4* (*I*) levels vary over time along the centriole surface, from 0 to 2π) subject to the initial conditions *A* = *A*_0_(1 + *W_A_*(*x*)) and *I* = *I*_0_(1 + *W_I_*(*x*)). Here, *A*_0_and *I*_0_are the homogeneous steady-state solutions to (3) and (4), and *W_A_* and *W_I_* are independent random variables with uniform distribution on [0, 1] that we use to generate the initial stochastic noise in the binding of *A* and *I* to the centriole surface at *t* = 0. All of the simulations that follow are performed over a unit of dimensionless time (*t* = 0 to *t* = 1), so the timescales of each simulation can be compared. All reaction and diffusion parameters in the system are large compared to unity, so all simulations achieve a steady-state within this unit of time. The dimensionless concentration values on the y-axis of the graphs shown in Figure 2B-D can be compared within these simulations.

We observe that the initial noise in the system is rapidly suppressed for *I*, and begins to be smoothed for *A* as the system approaches the steady state that would be stable in the absence of diffusion (Figure 2B, *t* = 0 to *t* = 0.01). However, on a slightly longer time-scale, this state is destabilised by the presence of diffusion (Figure 2B, *t* = 0.05) as any region in which the levels of *A* are slightly raised are reinforced due to the self-promotion of *A*. The same effect also increases *I*, but, since *I* rapidly diffuses away, the local accumulation of *A* is maintained while the production of *A* is inhibited around the rest of the centriole surface by the diffusing *I*. The solution therefore rapidly resolves to a non-homogeneous stable steady state with a single dominant peak (Figure 2B, *t* = 0.25). Both *A* and *I* continue to dynamically bind/unbind from the centriole in this steady state—in agreement with FRAP experiments showing that PLK4 continuously turns-over at centrioles (Cizmecioglu et al., 2010; Yamamoto and Kitagawa, 2019)—but this state is stable and remains unchanged for the remainder of the simulation (i.e. until *t* = 1; not shown). It is interesting to note that, in this model, it is mostly the activator species (i.e. non-phosphorylated, presumably kinase inactive, PLK4) that accumulates in the single PLK4 peak (Figure 2B), which may seem biologically implausible (see Discussion).

As described in the Introduction, it has been shown experimentally that overexpressing PLK4 leads to the generation of multiple PLK4 foci around the mother centriole whereas inhibiting PLK4 kinase activity prevents PLK4 symmetry breaking with PLK4 evenly distributed at high levels around the centriole surface. To test if Model 1 could recapitulate these behaviours, we simulated PLK4 overexpression by increasing the PLK4 production rate parameter *a* by 2-fold, and PLK4 kinase inhibition by reducing the phosphorylation rate parameter *b* by 5-fold. Doubling PLK4 production led to an increase in the number of transient PLK4 peaks that quickly settled to a stable steady-state of two peaks (Figure 2C). The two peaks were evenly spaced around the centriole, which is typical of Turing systems (see Discussion). In contrast, PLK4 symmetry was no longer broken when PLK4 kinase activity was reduced, and PLK4 accumulated evenly around the centriole to a high steady-state level (Figure 2D). This happens because less *I* (PLK4*) is produced when PLK4 kinase activity is reduced, so *I* can no longer suppress the accumulation of *A* (inactive PLK4) around the centriole. We conclude that, in this parameter regime, Model 1 can capture well three key features of PLK4 behaviour at the centriole: (1) Breaking symmetry to a single peak under appropriate conditions; (2) Breaking symmetry to more than one peak when PLK4 is overexpressed; (3) Failing to break symmetry and accumulating high levels of PLK4 when PLK4 kinase activity is inhibited.

#### Model 1 robustness analysis

To assess the robustness of Model 1’s ability to generate a single PLK4 peak when parameter values are changed, we generated a phase diagram showing the average number of PLK4 peaks generated over 20 simulations (shown in colour code) as we varied the rate of production of PLK4 (*a*) (the equivalent of varying PLK4s cytoplasmic concentration) and PLK4 kinase activity (*b*) (Figure 3A). Parameter values that do not support symmetry breaking are either indicated in *dark blue* (no PLK4 peaks, PLK4 distributed evenly at high levels around the centriole) or in *black* (no PLK4 peaks, no PLK4 present at the centriole). It can be seen that if PLK4 kinase activity drops below a certain level, the system robustly fails to break symmetry and PLK4 accumulates at high levels around the entire centriole surface (*dark blue* areas, Figure 3A). Thus, Model 1 robustly predicts the symmetric centriolar recruitment of high-levels of PLK4 when PLK4 kinase activity is inhibited (REFS).

**Figure 3.**
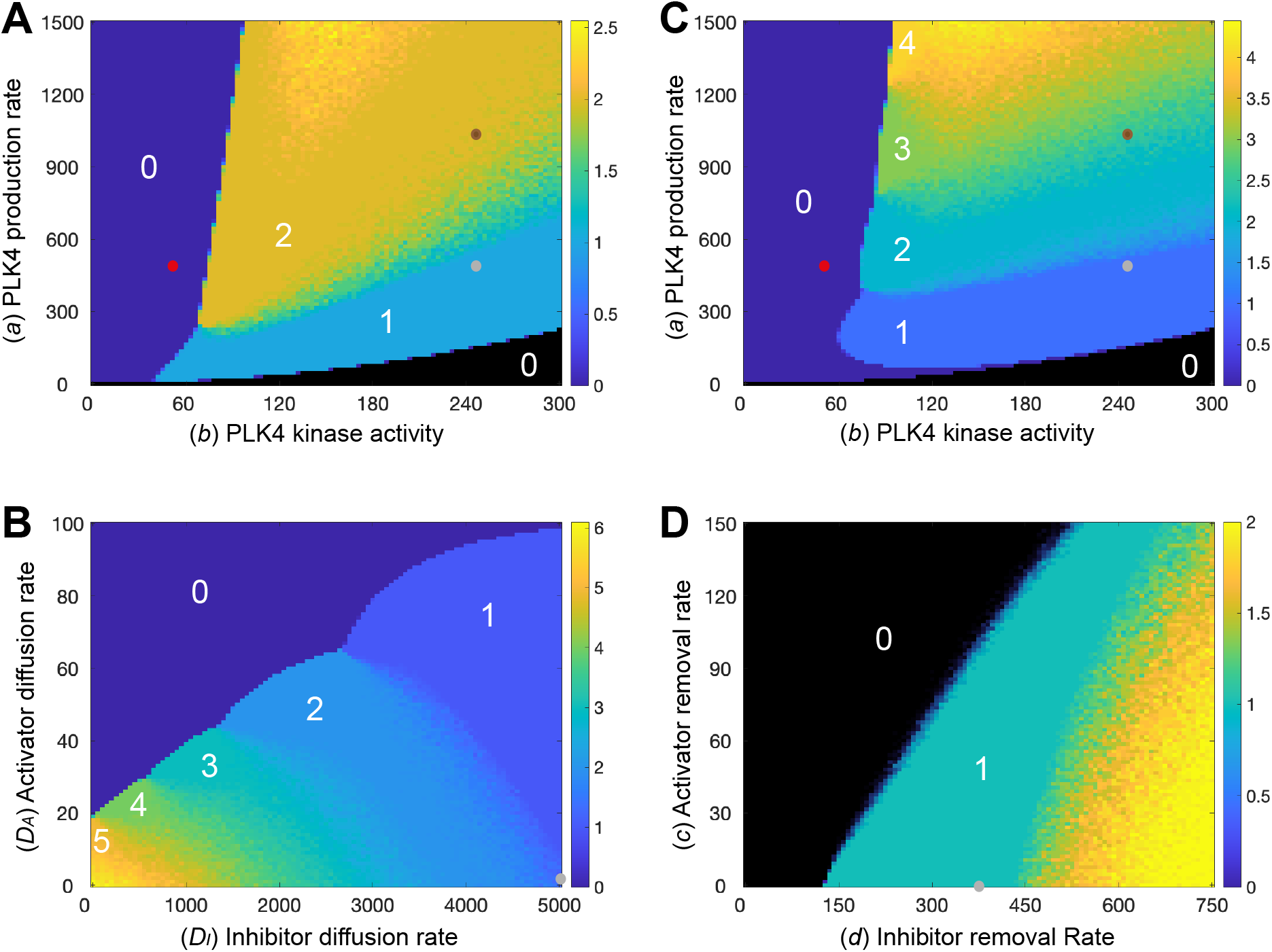
Analysis of the robustness of Model 1 to changes in parameter values. Phase diagrams show how the average number of PLK4 peaks generated around the centriole surface in 20 simulations (colour-coded to the scale shown on the right of each diagram) change as different parameters are varied: **(A,C)** The rate of PLK4 production (*a*) and PLK4 kinase activity (*b*) in Model 1 (A), or in a version of Model 1 in which we allow the diffusion rate of the PLK4 species to decrease as the levels of PLK4 in the system increase (C), see text for details; **(B)** The rate of diffusion of the activator (*D*_*A*_) and inhibitor (*D*_*I*_) species; **(D)** The rate at which the activator (*c*) or inhibitor (*d*) species are degraded/lost from the system. The number of peaks formed in certain phase spaces is highlighted (white numbers), and small dots indicate the parameter values used in the simulations shown in Figure 2: normal kinase levels and kinase activity (Figure 2B, *grey dots*), 2X PLK4 kinase levels (Figure 2C, *brown dots*) and 0.2X kinase activity (Figure 2D, *red dots*). Note that brown and red dots are not shown on (B,D) as kinase levels and activity remain constant in the simulations shown in these phase diagrams.

The phase diagram also reveals, however, that this system is not very robust at generating a single PLK4 peak as the amount of PLK4 is varied (i.e. the system cannot reliably produce a single PLK4 peak over a wide range of PLK4 concentrations). There are essentially no values of PLK4 production (*a*) that can reproducibly generate a single PLK4 peak (meaning that 20/20 simulations generated a single peak) that can still reproducibly generate a single PLK4 peak when *a* is either halved or doubled (Figure 3A, see Figure S1 for a more detailed illustration). Moreover, although increasing the production of PLK4 in this system generates more than one peak, it does not readily generate more than 2-3 peaks. This seems inconsistent with the biology, as the overexpression of PLK4 can lead to the assembly of up to 6 procentrioles around the mother centriole—see, for example, (Kleylein-Sohn et al., 2007; Vulprecht et al., 2012). It is not clear if these problems also occur in the original formulation of the Takao et al. model as no analysis of the robustness of the model to parameter changes was performed (Takao et al., 2019). As we discuss next, however, it seems likely that these are fundamental issues that will affect all models of this Type I system, leading us to propose a plausible solution.

In order for multiple peaks to occur in the overexpressed PLK4 limit, the diffusivity of the activator needs to be sufficiently small so that the multiple PLK4 peaks formed are thin enough to be accommodated around the centriole surface. An examination of the phase diagram comparing the number of PLK4 peaks formed when the diffusion rates of *A* (*D_A_*) and *I* (*D_I_*) are varied in Model 1 (Figure 3B) illustrates this point. We see that, as the diffusivities of *A* and *I* decrease, so the system forms increasing numbers of PLK4 peaks (up to 6 in the parameter regime analysed here). This is because, as their diffusivity decreases, *A* and *I* spread less efficiently around the centriole, allowing the formation of multiple, thinner, peaks. However, the decreasing diffusivity of *A* and *I* necessarily corresponds to a decreasing transfer of “information” between different regions of the centriole. Consequently, as the diffusivity of the PLK4 species decreases, it becomes increasingly difficult for the centriole to robustly form a single peak under normal conditions—since different regions of the centriole are not able to “communicate” with each other efficiently. In other words, single-peak robustness in one limit (low-levels of PLK4) and multiple peaks in another limit (high levels of PLK4) are fundamentally incompatible qualities of the system. Importantly, this issue is not limited to activator-inhibitor/diffusion-based models, but would apply to any spatial model in which information transfers around the centriole surface (as it presumably must do in the real-world physical system).

We realised that the well-characterised ability of PLK4 to dimerise to stimulate its own destruction via trans-autophosphorylation (Guderian et al., 2010; Holland et al., 2012; Cunha-Ferreira et al., 2013; Klebba et al., 2013) could potentially solve this problem. This is because any increase in cytoplasmic PLK4 levels will increase the probability of the PLK4 species generated at the centriole surface dimerising and degrading as they diffuse through the bulk cytoplasm. This means that any increase in the cytoplasmic levels of PLK4 will lead to a reduction in the effective diffusion of *A* and *I* around the centriole surface. If we modify Model 1 so that an increase in PLK4 production leads to a decrease in the diffusivity of the PLK4 species (see Appendix IV), the system now more robustly forms a single PLK4-peak in the low-PLK4 limit, while simultaneously forming a larger number of peaks in the high-PLK4 limit (Figure 3C; see Figure S1 for a more detailed analysis). In a more general sense, we propose that increasing cytoplasmic PLK4 levels slows down the transfer of information around the centriole due to destructive dimerization; this mechanism permits increasing levels of overduplication in the high-PLK4 limit without affecting system robustness under normal conditions.

Finally, to analyse the effect of varying the dissociation/degradation rates of *A* (*c*) and *I* (*d*) we also generated a (*c*, *d*) phase diagram (Figure 3D). We find that the system is able to break symmetry provided that (*c*) is below a certain threshold that depends on (*d*). If (*c*) is above this threshold, then *A* becomes fully depleted from the system, which in turn removes the source of *I* (*black region*, Figure 3D)

### Model 2: Inhibitor to Activator conversion based on the model of Leda et al., 2018

The second model we analyse is a Type II system in which a rapidly-diffusing inhibitor (*I*) is converted into a slowly-diffusing activator (*A*) through phosphorylation. As a biological example of such a model, we adapt the reaction regime proposed by Leda et al (2018). This system comprises just two proteins, PLK4 and STIL, but these combine to create four components defined by the phosphorylation state of each component, [*PS*], [*P*^∗^*S*], [*PS*^∗^], and [*P*^∗^*S*^∗^] (* denoting phosphorylation). The authors assume that each component can bind to the centriole surface and may be converted into another, either in the cytoplasm or at the centriole surface, through phosphorylation/dephosphorylation, with the [*P*^∗^*S*^∗^] species promoting the phosphorylation of all the other species. The centriole complexes in which STIL is phosphorylated are postulated to exchange with the cytoplasm at a slower rate than the complexes in which STIL is not phosphorylated (this will be the basis for the difference in the diffusion rates of the two-component system we formulate below). As in the Takao et al. model, Leda et al. allow the complexes to bind to discrete centriole compartments, but, importantly, there is no ordering or spatial orientation of these compartments. Instead, compartmental exchange occurs through the shared cytoplasm.

By reinterpreting this cytoplasmic exchange as a diffusive process—with diffusivity dependent on the rate of exchange—we develop a Turing system based on the reactions of the Leda et al. model (Model 2; Figure 4A). The two relevant components in the system are not PLK4 and STIL, but rather PLK4:STIL complexes that act as either fast-diffusing Inhibitors (*I*) comprising PLK4 bound to non-phosphorylated STIL ([*PS*] and [*P*^∗^*S*]), or slow-diffusing Activators (*A*) comprising PLK4 bound to phosphorylated STIL ([*PS*^∗^] and [*P*^∗^*S*^∗^]). The derivation of this model is given in detail in Appendix III which leads to the system:

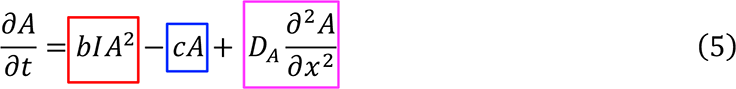

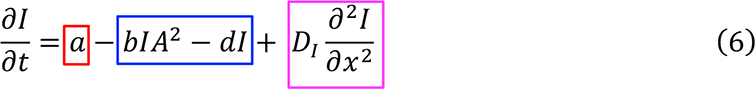

**Figure 4.**
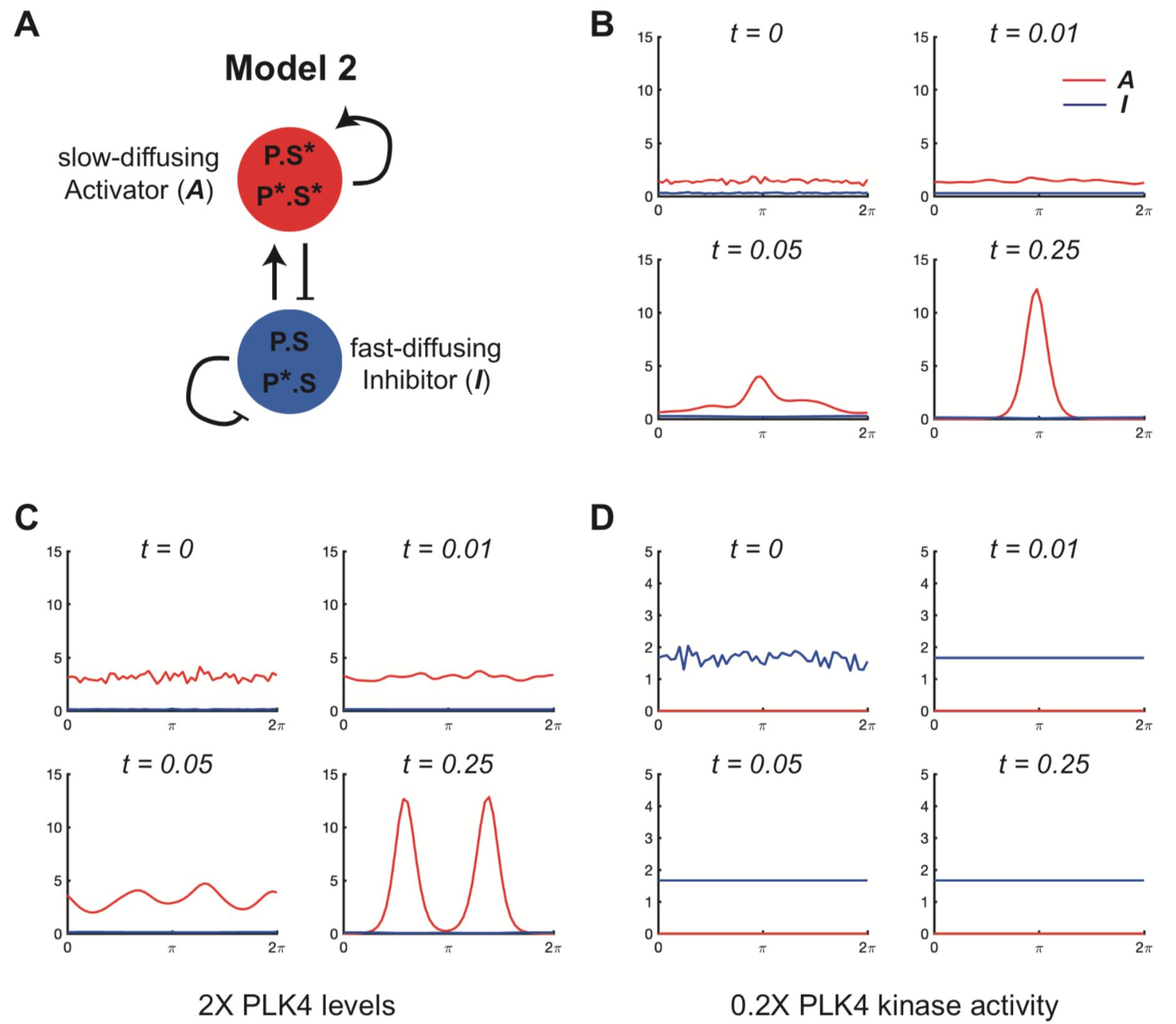
Computer simulations of mathematical Model 2. **(A)** Schematic summarises the reaction regime of Model 2, a Type II Turing-system. The biology underlying this model was originally proposed by Leda et al., (2018). Although it is not possible to precisely assign the activator and inhibitor relationships depicted in this schematic to specific terms in the reaction equations (equations 5 and 6, see main text) we can approximately see that *A* (which contains the active P*S* species that phosphorylates all other species) simultaneously generates more *A* and inhibits *I* by converting nearby species that contain non-phosphorylated STIL (*I*) into complexes containing phosphorylated STIL (*A*). *I* promotes the formation of *A* because it acts as the source of *A* in the conversion process, and *I* is self-inhibiting through promoting its own dissociation/degradation (i.e. the rate of dissociation/degradation of *I* is proportional to the amount of *I* – eq.6). **(B-D)** Graphs show how the levels of the activator species (*red* lines) and the inhibitor species (*blue* lines) change over time (arbitrary units) at the centriole surface (modelled as a continuous ring, stretched out as a line between 0-2ν) in computer simulations. The rate parameters for each simulation are defined in the main text and can lead to symmetry breaking to a single peak (B), or to multiple peaks when cytoplasmic PLK4 levels are increased (C), or to no symmetry breaking at all—with the near-complete depletion of the activator species from the centriole surface, and the uniform low-level accumulation of the inhibitor species—when PLK4 kinase activity is reduced (D).

As before, the boxed parts of the equation describe how *A* (eq. 5) or *I* (eq. 6) are produced (*red*), removed (*blue*) or diffuse (*magenta*) within the system. Here, *a* is a constant source term for the production of *I* (i.e. the rate at which phosphorylated and non-phosphorylated PLK4 molecules form complexes with STIL), *b* is the rate at which *I* is converted into *A*, (i.e. the rate at which STIL is phosphorylated within these complexes), and *c* and *d* are the rates at which the *A* and *I* species, respectively, are degraded or lost to the cytoplasm.

As mentioned above, it is not possible to precisely assign the activator and inhibitor relationships depicted in the schematic in Figure 4A to specific terms in the reaction equations (eq.5 and eq.6), but an approximate explanation is provided in the Figure legend. The conditions for this system to break symmetry are derived in Appendix II and, unless stated otherwise, we choose the parameter values *a* = 100, *b* = 150, *c* = 60, *d* = 60, *D_A_* = 2, and *D_I_* = 5000, that satisfy these conditions. These values were chosen in part to reflect the parameter values and ratios used in the previous modelling papers (Leda et al., 2018; Takao et al., 2019).

In Figure 4B, we plot the solution output subject to the initial conditions *A* = *A*_0_(1 + *W_A_*(*x*)) and *I* = *A_I_*(1 + *W_I_*(*x*)), where *A*_0_ and *I*_0_are the homogeneous steady-state solutions to (5) and (6) and *W_A_* and *W_I_* are independent random variables with uniform distribution on [0, 1]. As with Model 1, all of the simulations that follow are performed over a unit of dimensionless time and all reaction and diffusion parameters in the system are large compared to unity, so all simulations achieve a steady state within this unit of time. The dimensionless concentration values on the y-axis of the graphs shown in Figure 4B-D can be compared to each other, but not to the values shown in Figure 2B-D (for Model 1) as these dimensionless values depend on dimensional reaction rates, which differ between the two models.

As with Model 1, the solution approaches a stable non-homogeneous steady state with a dominant peak after an initial smoothing period. In contrast to Model 1, we observe that, in the region of the activator peak, the inhibitor exhibits a slight dip. This is because the activator promotes the production of the inhibitor in Model 1, but suppresses the production of the inhibitor (by promoting its conversion to the activator) in Model 2. We then simulated PLK4 overexpression in the system by increasing the production rate parameter of the PLK4:STIL complexes (*a*) by 2-fold (Figure 4C), and PLK4 kinase inhibition by reducing the phosphorylation rate parameter (*b*) by 5-fold (Figure 4D). As with Model 1, doubling PLK4:STIL production led to an increase in the number of transient PLK4 peaks that quickly settled to a stable steady-state of two peaks that were evenly spaced around the centriole. Interestingly, although inhibiting PLK4 kinase activity led to a failure to break symmetry, *A* was no longer detectable at the centriole and *I* accumulated only at a low uniform level. Thus, unlike Model 1, Model 2 does not capture well the high-level accumulation of kinase-inhibited PLK4 species that has been observed experimentally.

#### Model 2 robustness analysis

To assess the robustness of Model 2 to changes in parameter values we first generated a phase-diagram showing the average number of PLK4 peaks generated over 20 simulations as we varied the rate of PLK4:STIL production (*a*) and PLK4 kinase activity (*b*) (Figure 3A). Parameter values that do not support symmetry breaking are indicated in *dark blue.* In the limit of high PLK4 levels and high kinase activity, the system accumulates high levels of activator uniformly around the centriole (*dark blue* region, top-right of Figure 3B). This is analogous to all compartments being occupied in a discrete model, and would likely result in multiple daughter centrioles being produced. By contrast, at low levels of kinase activity, there is a low-level uniform distribution of the inhibitor, and no accumulation of the activator (*dark blue* region, bottom left of Figure 3B). Therefore, Model 2 robustly fails to recapitulate the high-level centriolar accumulation of PLK4 observed when PLK4 kinase activity is inhibited (Ohta et al., 2018; Yamamoto and Kitagawa, 2019).

Moreover, in this parameter regime, Model 2 suffers from the same two problems we encountered with Model 1: (1) There is no value of *a* that can robustly generate a single peak of PLK4 that can still do this when (*a*) is halved or doubled (Figure S1); (2) The system struggles to generate >3-4 PLK4 peaks when PLK4 is overexpressed. As before, decreasing the diffusivity of *A* and *I* allows the system to generate more, thinner, peaks (Figure 5B), and linking the increase in PLK4 production to a decrease in diffusivity (as we did for Model 1 – see Appendix IV) allows the system to generate more (>6) centrioles when PLK4 is overexpressed, although it has a less pronounced effect on the robustness of the system to produce a single PLK4 peak (Figure 5C; Figure S1). Interestingly, the original Leda et al. model (Leda et al., 2018) avoids these problems because it supposes no spatial relationship between the individual compartments and instead assumes that communication between compartments is instantaneous. However, this requires that diffusion is sufficiently fast that concentration gradients are negligible between centriolar compartments, but not so fast that the relevant species are diluted in the cytoplasm. It seems implausible that both of these effects could be achieved with a single diffusion rate in a real-world physical system where the centrioles exist in the context of a much larger volume of cytoplasm.

**Figure 5.**
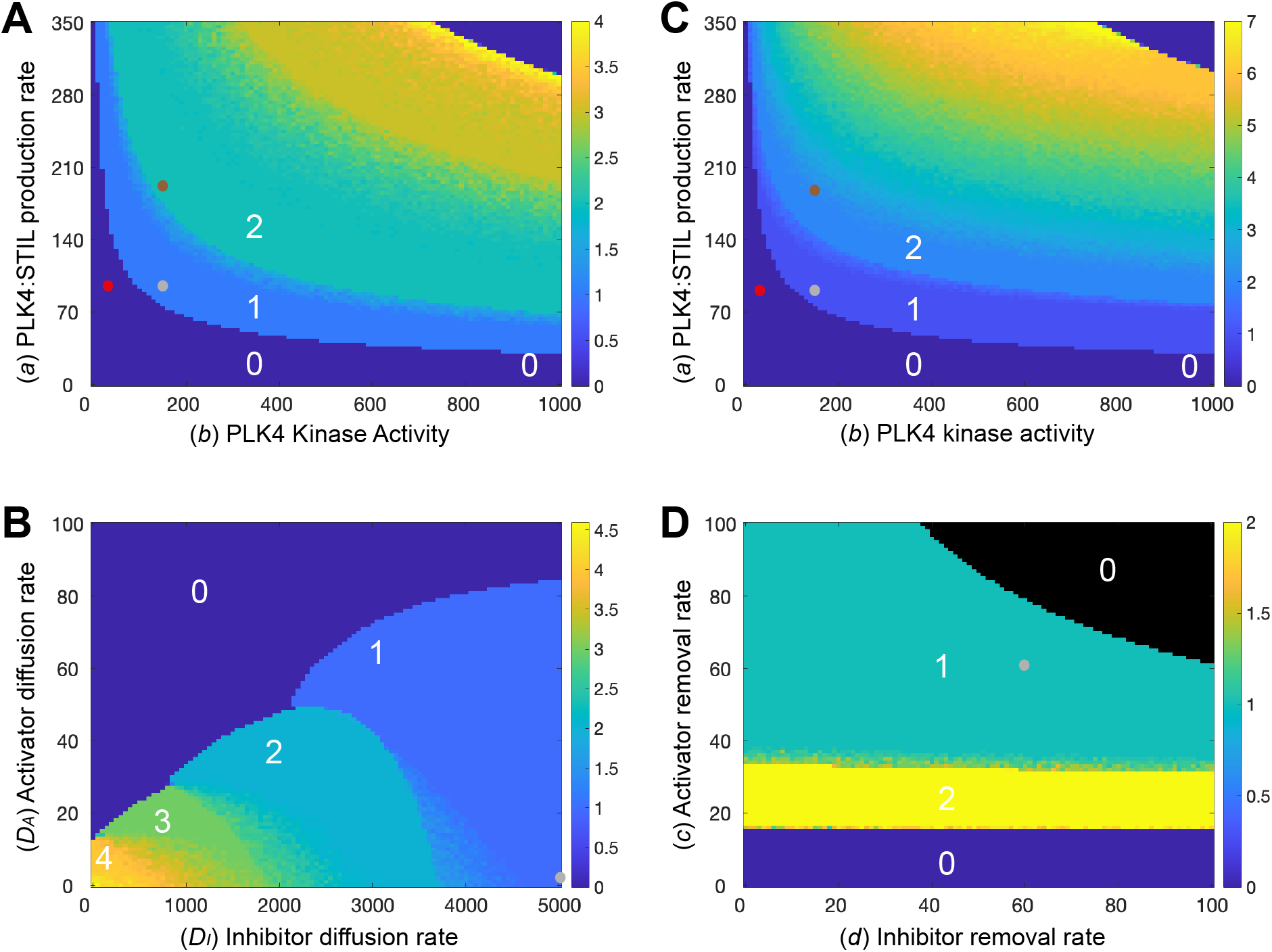
Analysis of the robustness of Model 2 to changes in parameter values. Phase diagrams show how the average number of PLK4 peaks generated around the centriole surface in 20 simulations (colour-coded to the scale shown on the right of each diagram) change as different parameters are varied: **(A,C)** The rate of PLK4 production (*a*) and PLK4 kinase activity (*b*) in Model 2 (A), or in a version of Model 2 in which we allow the diffusion rate of the PLK4 species to decrease as the levels of PLK4 in the system increase (C), see text for details; **(B)** The rate of diffusion of the activator (*D*_*A*_) and inhibitor (*D*_*I*_) species; **(D)** The rate at which the activator (*c*) or inhibitor (*d*) species are degraded/lost from the system. The number of peaks formed in certain phase spaces is highlighted (white numbers), and small dots indicate the parameter values used in the simulations shown in Figure 4: normal kinase levels and kinase activity (Figure 4B, *grey dots*), 2X PLK4 kinase levels (Figure 4C, *brown dots*) and 0.2X kinase activity (Figure 4D, *red dots*). Note that brown and red dots are not shown on (B,D) as kinase levels and activity remain constant in the simulations shown in these phase diagrams.

The (*c*, *d*) phase diagram (Figure 5D) shows that, provided (*c*) is above a certain threshold, there is a large region of the parameter space in which the system breaks symmetry to a single peak. For values of (*c*) below this threshold, the species accumulate uniformly around the centriole due to the low degradation rate. In contrast, for sufficiently large values of (*c*) and (*d*), both species are fully depleted from the system.

### Unifying the models of Takao et al., and Leda et al

At a first glance, the original models of Takao et al. and Leda et al. appear to be very different: they use different mathematical methods to describe different chemical reactions, with symmetry breaking in the former being driven by PLK4 phosphorylation and in the latter by the phosphorylation of STIL in complexes with PLK4. In deriving Model 2, we grouped together the species *A* = [*PS*^∗^] + [*P*^∗^*S*^∗^] and *I* = [*PS*] + [*P*^∗^*S*] based on the unbinding rates of the four species specified in the Leda et al. paper—where complexes containing phosphorylated STIL ([*PS*^∗^] and [*P*^∗^*S*^∗^]) unbind slowly and those containing non-phosphorylated STIL ([*PS*] and [*P*^∗^*S*]) unbind rapidly. It is simple, however, to modify these model parameters so that the unbinding rate now depends on the phosphorylation state of PLK4 in these complexes (as is the case in the Takao et al. model), rather than on the phosphorylation state of STIL—i.e. we allow [*P*^∗^*S*] and [*P*^∗^*S*^∗^] to now unbind rapidly and [*PS*] and [*PS*^∗^] to now unbind slowly. Thus, in this reinterpretation, we are essentially applying the biological justification of the Takao et al. model to the Leda et al. model.

In this scenario, if we set *A* = [*PS*] + [*PS*^∗^] and *I* = [*P*^∗^*S*] + [*P*^∗^*S*^∗^] then, following the same procedure as before (see Appendix III), we arrive at a new model,

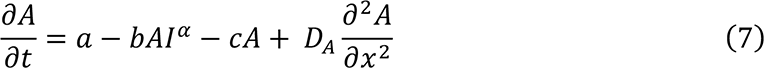

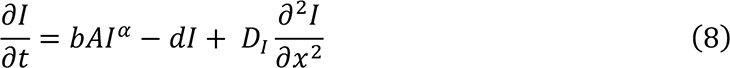

We observe that by setting *α* = 1/2 and substituting the sigmoidal self-assembly source term *aA*^2^/(1 + *A*) in place of the constant source term, *a*, we obtain *exactly* Model 1. In other words, all of the dynamics of the Takao et al. model is contained within the Leda et al. model, but with additional complexity and a different choice of rate parameters.

### Modelling symmetry breaking on a compartmentalised centriole surface

In our modelling so far, PLK4 symmetry breaking occurs on a continuous centriole surface; it does not require that the centriole surface be divided into discrete compartments that effectively compete with each other to become the dominant site (as is assumed in the previous models). Importantly, however, our model still applies if we divide the centriole surface into an arbitrary number of discrete compartments, with the various PLK4 species interacting only within an individual compartment, but diffusing between the spatially separated compartments laterally (see Appendix V). In Figure 6, we show the system outputs of Model 1 (Figure 6A) and Model 2 (Figure 6B) when we run simulations with a centriole surface comprising 9 discrete compartments that interact with the various PLK4 species. In both instances, the systems robustly break their initial symmetry to produce a single dominant compartment that accumulates the relevant species. This result will hold for an arbitrary number of compartments, as shown in Figure S2A (Model 1) and Figure S2B (Model 2), for compartment numbers 2 – 10.

**Figure 6.**
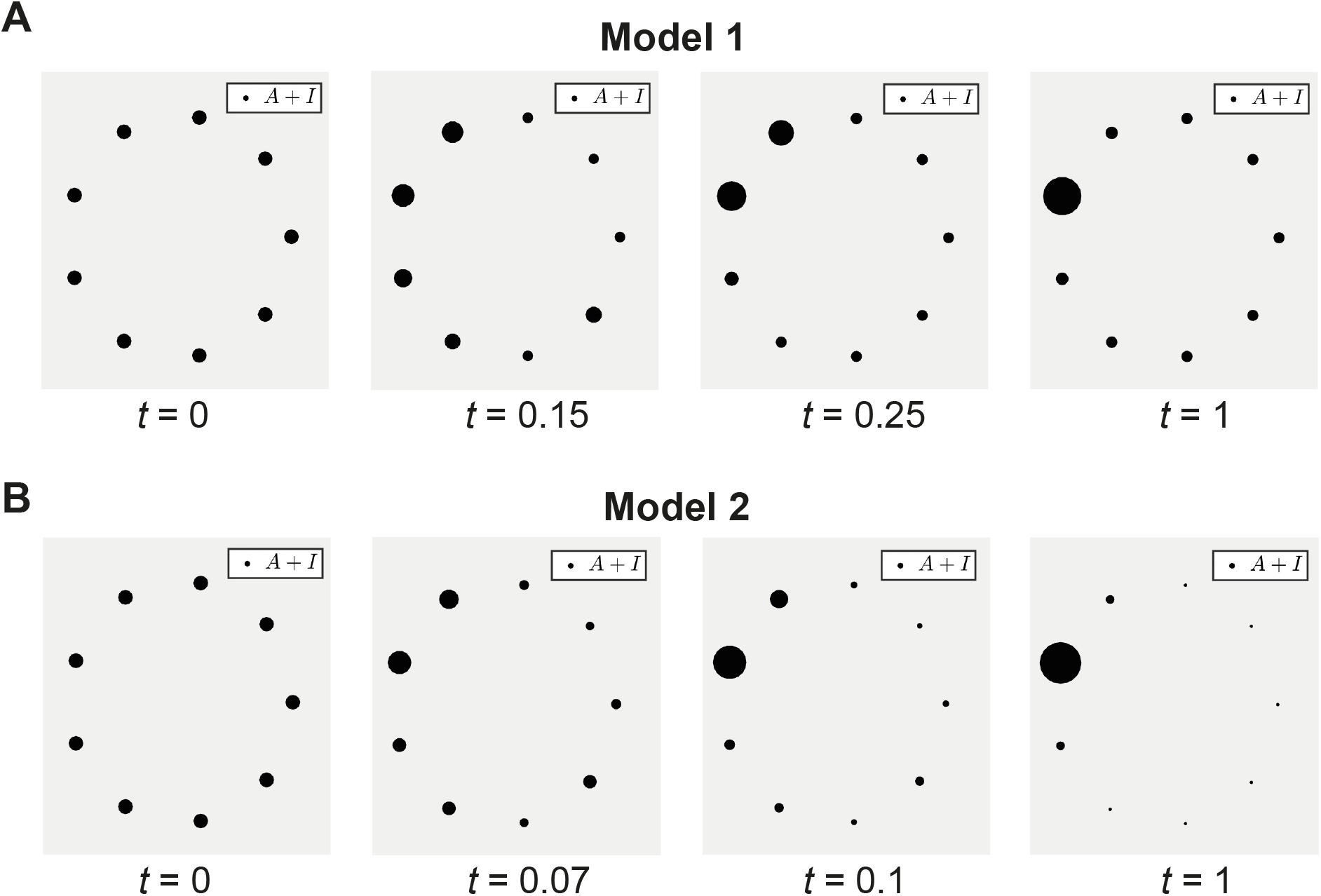
Analysis of PLK4 symmetry breaking on a 9-fold symmetric centriole surface. Diagrams show the output of computer simulations of Model 1 **(A)** and Model 2 **(B)** that have been adjusted so that the centriole is no longer one continuous surface but rather 9 independent compartments. Each compartment is depicted here as an individual dot distributed equally around the circumference of the centriole, with the size of each dot representing the amount of activator and inhibitor present in that compartment. At *t* = 0, the activator and inhibitor species are allowed to bind to and diffuse between the 9 different compartments. In both models, the system rapidly breaks symmetry, initially exhibiting a considerable variation in the amount of *A* and *I* recruited to each compartment. Both systems then settle to a dynamic steady state with a single dominant peak.

## Discussion

PLK4 is the master regulator of centriole biogenesis, so understanding how it becomes concentrated at a single site on the side of the mother centriole is crucial to understanding how centriole numbers are so strictly controlled (Fırat-Karalar and Stearns, 2014; Loncarek and Bettencourt-Dias, 2018; Nigg and Holland, 2018; Breslow and Holland, 2019). Here, we mathematically modelled PLK4 symmetry breaking as a simple two-component “Turing system”. All such models can be classified as either Type I or Type II systems depending on their reaction dynamics (Gierer and Meinhardt, 1972; Ruan, 1998) (Figure 1C), and we presented a Type I model based on the reaction scheme originally proposed by Takao et al. (2019), and a Type II model based on the reaction scheme originally proposed by Leda et al (2018). We note that Takao et al. described their system as being analogous to a “Turing model”, but it is not a reaction-diffusion model, and it does not exhibit the property of long-range inhibition that is central to all Turing-systems to produce a single PLK4 peak. Instead, it uses lateral inhibition (in which the influence of the inhibiting species does not extend beyond the neighbouring compartments) to reduce the number of potential PLK4 binding sites from ∼12 to ∼6. A single winning site is subsequently selected when STIL is added to the system— with additional positive feedback (not involving reaction-diffusion) ensuring that the compartment with most PLK4 becomes the dominant site.

Although these previous models used different mathematical methods and proposed different reaction regimes, we find that both models may be described by the same Turing-system dynamics. Thus, although it was not appreciated at the time, both previous attempts to model PLK4 symmetry breaking have essentially arrived at the same solution—with phosphorylated and non-phosphorylated species containing PLK4 (either on its own or in a complex with STIL) acting as either a slow diffusing activator (*A*) or a fast-diffusing inhibitor (*I*) species—with the differential centriole binding and/or unbinding properties of these species effectively allowing them to diffuse around the centriole at different rates.

Importantly, it can be mathematically proven that all such Turing-systems that break symmetry must satisfy either the Type I or Type II criteria, and that there are well-defined mathematical constraints on the parameter values required to support symmetry breaking in these systems. Nevertheless, we suspect that it will be challenging to precisely identify the molecular species that comprise *A* and *I*, and to precisely define the chemical relationship between them. This is because the interactions of PLK4 with itself (Park et al., 2014; Shimanovskaya et al., 2014; Slevin et al., 2012; Cottee et al., 2017; Leung et al., 2002; Arquint et al., 2015), with its centriole receptors CEP152/Asl and CEP192/Spd-2 (Hatch et al., 2010; Dzhindzhev et al., 2010; Cizmecioglu et al., 2010; Sonnen et al., 2013; Kim et al., 2013; Park et al., 2014; Boese et al., 2018) and with other crucial centriole duplication proteins (such as STIL/Ana2) (Moyer et al., 2015; Dzhindzhev et al., 2017; Ohta et al., 2018; McLamarrah et al., 2018; Moyer and Holland, 2019; McLamarrah et al., 2020) are complex. PLK4 therefore probably binds to the centriole surface through a web of interactions involving several species. It is for this reason that we have not attempted here to make extensive predictions about the molecular identity of *A* and *I* or their precise reaction regimes. Nevertheless, by probing these interactions in a general setting, it may be possible to establish several properties that *A* and *I* must satisfy without explicitly knowing their composition, which in turn may help in identifying them. We highlight 3 questions below that illustrate how our modelling can inform future thinking about this problem.

### 1. Could kinase-inactive PLK4 accumulate at the site of daughter centriole assembly?

Our modelling makes it clear that there are potential problems with the biology underlying both previous models of PLK4 symmetry breaking. In models in which phosphorylated species of PLK4 act as fast-diffusing inhibitors (as in Model 1, based on Takao et al. 2019) it is the non-phosphorylated (and so presumably kinase inactive) PLK4 species that will usually accumulate to the highest levels within the PLK4 peak (Figure 2B). *A priori*, this seems implausible, as PLK4 is thought to phosphorylate STIL to drive centriole assembly (Ohta et al., 2014; Dzhindzhev et al., 2014; Kratz et al., 2015). On the other hand, in models in which phosphorylated species of PLK4 act as slow diffusing activators (as in Model 2, based on Leda et al., 2018), inhibiting PLK4 kinase activity will lead to the low-level accumulation of the faster turning-over non-phosphorylated species and the complete loss of the slowly turning over phosphorylated species (Figure 4D). This is inconsistent with data showing that inhibiting PLK4 kinase activity leads to the high-level accumulation of slowly turning-over PLK4 (Yamamoto and Kitagawa, 2019). How can we resolve these issues?

Given the increasing evidence that inhibiting PLK4 kinase activity leads to the high-level accumulation of slowly turning-over PLK4, which is clearly inconsistent with Model 2, we think it is worth considering the possibility that it is the inactive form of PLK4 that normally accumulates in the PLK4 peak (as predicted by Model 1). Perhaps this doesn’t matter, as in Type I systems some active kinase will also accumulate in this peak (Figure 2B). This small amount of active kinase may be sufficient if, for example, inactive PLK4 is primarily responsible for recruiting proteins such as STIL/Ana2 and SAS6 to this site, and only a small amount of active PLK4 kinase is required to stimulate daughter centriole assembly. An alternative possibility is that PLK4 kinase activity is only required for PLK4 symmetry breaking, and not daughter centriole assembly. There is some evidence to support this possibility, as in some systems PLK4 kinase activity appears to only be required at the end of mitosis/early G1—when PLK4 symmetry is presumably being broken, but before daughter centrioles have physically started to assemble (Zitouni et al., 2016). Moreover, when co-overexpressed in early *Drosophila* embryos, Sas-6 and STIL/Ana2 assemble into large Particles (SAPs) that recruit many other centriole proteins (Stevens et al., 2010; Gartenmann et al., 2020). SAP assembly does not require PLK4, yet it appears to be stimulated by the phosphorylation of STIL/Ana2 (Gartenmann et al., 2020), suggesting that another kinase may phosphorylate STIL/Ana2 to promote daughter centriole assembly.

### 2. How are multiple PLK4 peaks spaced around the centriole surface?

A feature of Turing-systems is that any multiple peaks formed will become evenly spaced within the system if given enough time to do so (as peaks are most stable when they are as far apart from each other as possible). Thus, a prediction of our modelling is that multiple PLK4 peaks will ultimately become evenly spaced around the centriole surface. Unfortunately, the spacing of multiple PLK4 peaks in cells overexpressing PLK4 has not been quantified. Thus, one is left to interpret published images, some of which might support equal spacing while others appear not to (e.g. Kleylein-Sohn et al, Dev. Cell, 2007). Moreover, this analysis is complicated because CEP152/Asl, a major centriolar recruiter of PLK4 (Hatch et al., 2010; Dzhindzhev et al., 2010; Cizmecioglu et al., 2010), can form incomplete rings (Andrew Holland and Jadranka Loncarek, *pers. comm.*). This can be appreciated when considering Figure 2C in (Hatch et al., 2010), where the extra centrioles induced by PLK4 overexpression are not evenly spaced around the centriole, but are approximately evenly spaced around the incomplete CEP152 ring.

### 3. Are PLK4 receptors organised into discrete compartments?

A unique benefit of our modelling approach is that it does not rely on PLK4 being recruited to centrioles by receptors that are organised into discrete compartments, as was the case in both previous models. While there is some data supporting the idea that centriolar CEP152/Asl, a major recruiter of PLK4 to the centriole, may be organised into discrete compartments, the number and organisation of these compartments is unclear (Takao et al., 2019; Tian et al., 2021; Wainman, 2021; Gao et al., 2021; Tian et al., 2022; Wilmerding et al., 2023). Moreover, it has been proposed that CEP152 self-assembles into a continuous ring around the centriole (Kim et al., 2019; Lee et al., 2020), while the most recent super-resolution studies suggest that CEP152 is not normally detectably compartmentalised around the centriole surface (Jadranka Loncarek and Andrew Holland*, pers. comm.*). Interestingly, however, if PLK4 kinase activity is inhibited, PLK4 and CEP152 coalesce into up to 9 discrete compartments over several (∼24) hours (Andrew Holland and Jadranka Loncarek, *pers. comm.*). This coalescence presumably occurs through a process that is distinct from normal PLK4 symmetry breaking, which requires PLK4 activity and occurs on a faster timescale.

In summary, our modelling approach reveals that PLK4 symmetry breaking can be explained using the simple and well-studied Turing-system framework that readily permits analysis and comparison. This approach allows us, for the first time, to start to understand why these systems can break symmetry. We propose that the underlying biological process driving PLK4 symmetry breaking is a diffusion-driven instability caused by short-range activation and long-range inhibition.

---

**Figure S1.**
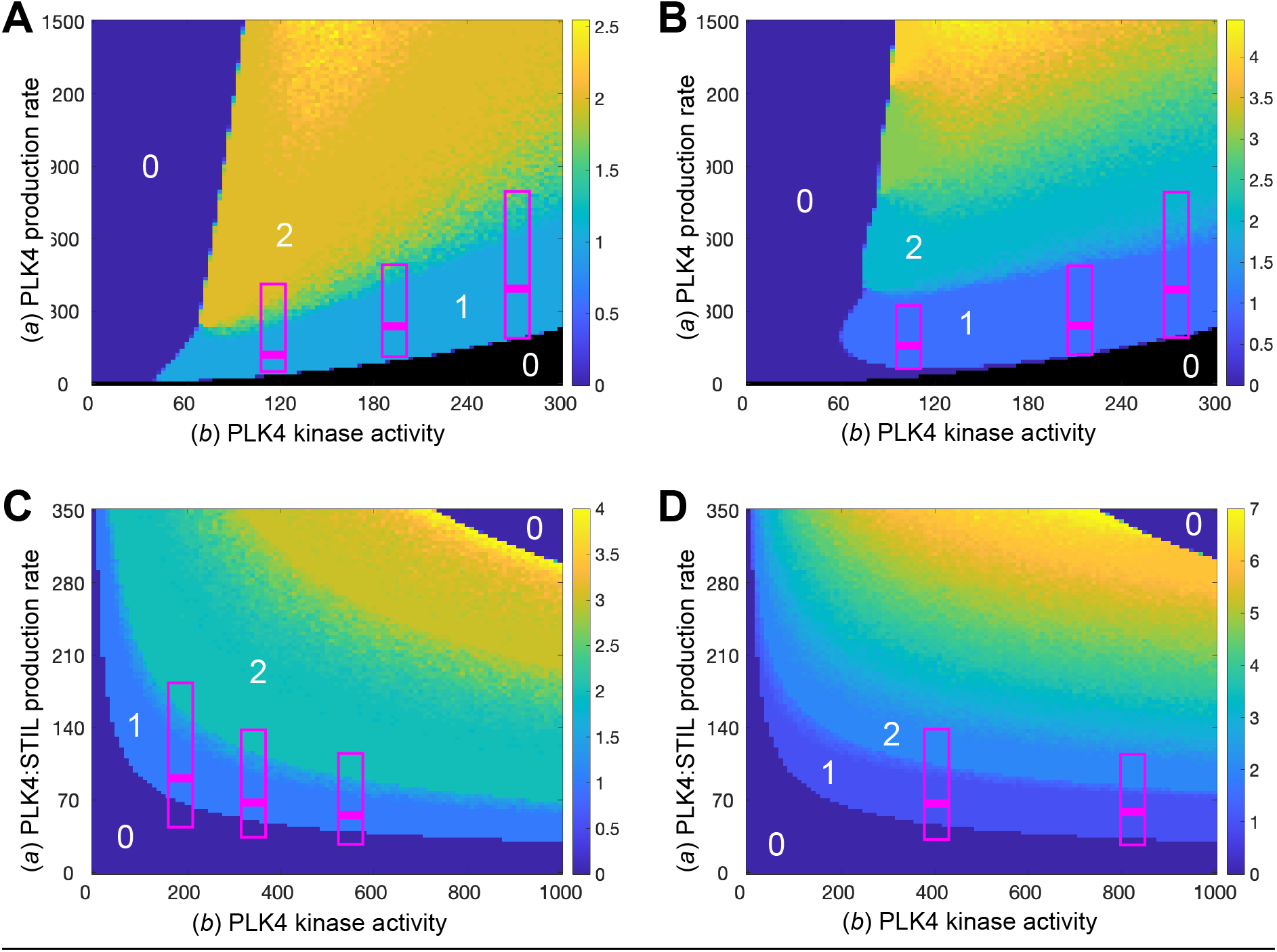
Analysing system robustness if the diffusion rate of the activator and inhibitor species is allowed to decrease as the concentration of PLK4 increases. **(A)** The phase diagram shown here is the same as that shown in Figure 3A comparing Model 1’s robustness to changes in the rate of PLK4 production and kinase activity. The *magenta* boxes each highlight a particular concentration of PLK4 (thick horizontal line) and the vertical height of the boxes show the values when that concentration is halved (bottom of boxes) or doubled (top of boxes). It can be seen that in this original version of Model 1 there is essentially no concentration of PLK4 that can robustly generate a single PLK4 peak (*light blue* area) that can still robustly generate a single PLK4 peak when its concentration is either halved and doubled. Also, when these concentrations of PLK4 are increased beyond doubling, they do not readily generate more than 2-3 PLK4 peaks. **(B)** The phase diagram shown here is the same as that shown in Figure 3C, where we adapted Model 1 to allow the diffusivity of the PLK4 species to decrease as the concentration of PLK4 increases. In this adapted model it can be seen that the concentrations of PLK4 highlighted with the *magenta* horizontal lines and associated boxes can still robustly generate a single PLK4 peak when that concentration of PLK4 is either halved or doubled, and would also more readily form multiple peaks of PLK4 when PLK4 is overexpressed to even higher levels. **(C,D)** Phase diagrams show the same analysis of Model 2 as described in (A,B) above for Model 1.

**Figure S2.**
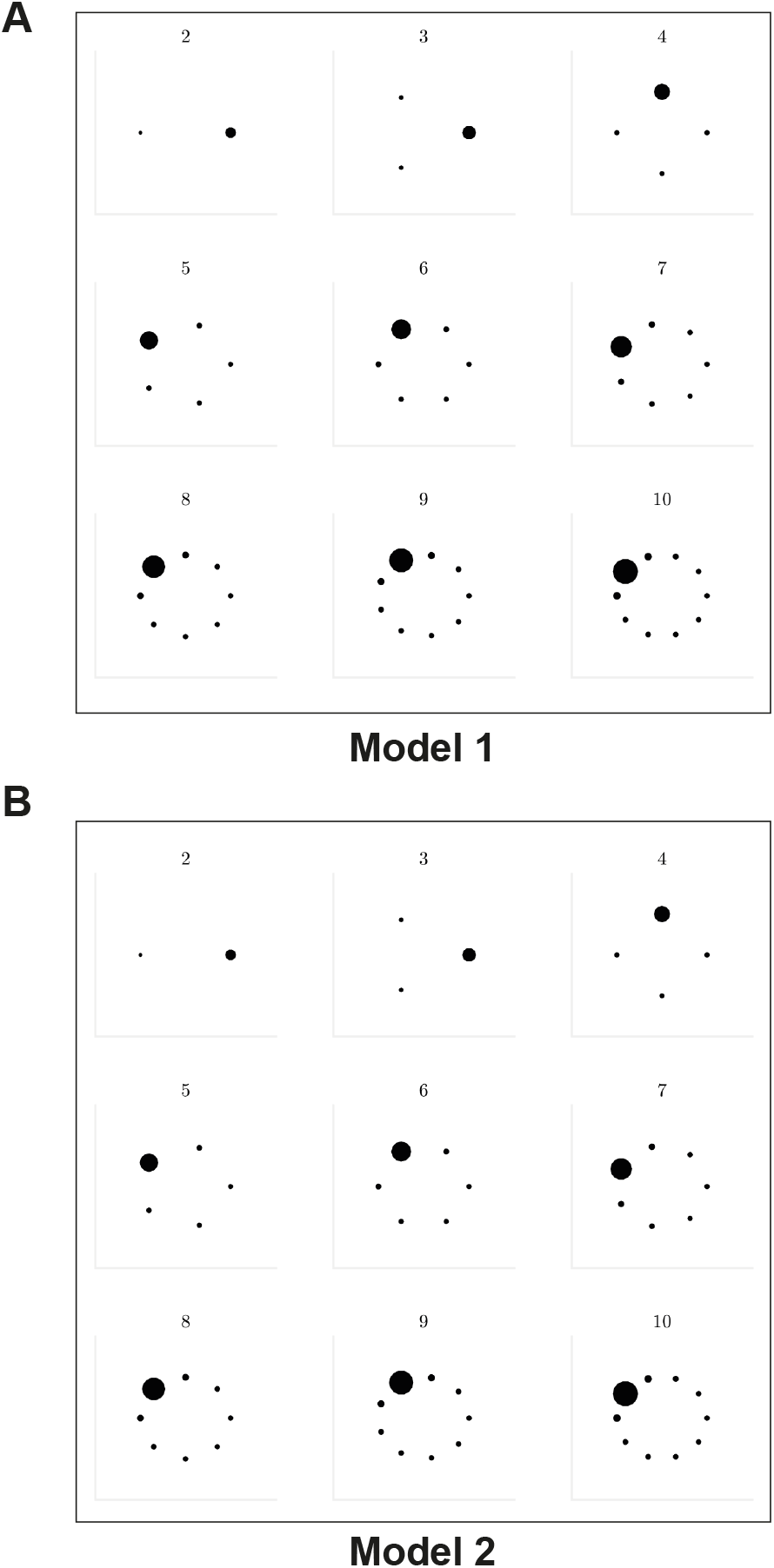
Analysis of PLK4 symmetry breaking on a centriole surface comprising different numbers of PLK-binding compartments. Diagrams show the output of computer simulations of Model 1 **(A)** and Model 2 **(B)** that have been adjusted so that the centriole is no longer one continuous surface but rather 2-10 independent compartments. Each compartment is depicted here as an individual dot distributed equally around the circumference of the centriole, with the size of each dot representing the amount of activator and inhibitor present in that compartment at the end of the simulation. Both systems settle to a dynamic steady state with a single dominant peak no matter how many compartments are present around the centriole surface.

