## Appendix I-V for "A simple Turing reaction-diffusion model can explain how mother centrioles break symmetry to generate a single daughter"

### Appendix I: Conditions for symmetry-breaking in a Turing-system

In order to break symmetry, there are two conditions that a two-species Turing reaction-diffusion model must satisfy at the steady state (Ruan, 1998): (a) the steady state must be stable in the absence of diffusion; (b) this steady state must be unstable in the presence of diffusion. In order to satisfy conditions (a) and (b), several mathematical inequalities must be satisfied. First, we require that precisely one of the two species be self-promoting and the other be self-inhibiting. Mathematically, starting from the general form of the coupled reaction-diffusion equations

$$\frac{\partial A}{\partial t} = f(A, I) + D_A \frac{\partial^2 A}{\partial x^2}\,, \tag{A1}$$

$$\frac{\partial I}{\partial t} = g(A, I) + D_I \frac{\partial^2 I}{\partial x^2}\,, \tag{A2}$$

these two conditions (self-promotion and self-inhibition) imply that

$$\frac{\partial f}{\partial A} > 0, \quad \frac{\partial g}{\partial I} < 0 \tag{A3}$$

where these inequalities, and all that follow, are evaluated at the steady state that we wish to analyse for its symmetry-breaking properties. Indeed, since $f$ is the rate at which $A$ is produced, $\frac{\partial f}{\partial A} > 0$ implies that an increase in the concentration of $A$ leads to an increase in $f$ and therefore to an increase in the rate of production of $A$. Hence, this constraint implies that $A$ is self-*promoting*. Conversely, by the same reasoning, $\frac{\partial g}{\partial I} < 0$ implies that $I$ is self-*inhibiting.* Negative gradients in this context correspond to stabilising effects whereas positive gradient correspond to destabilising effects. These inequalities provide a mathematical distinction between activator and inhibitor. If both species were to be self-promoting then any positive perturbation to the steady state would cause both species to continuously accumulate. In this case, the system would be unstable even in the absence of diffusion; this contradicts condition (a). On the other hand, if both species were self-inhibiting then any positive perturbation to the system would decay and the steady state would be stable, even in the presence of diffusion; this contradicts condition (b). As an additional constraint, we also require that the stabilising strength of the inhibitor be greater than the destabilising strength of the activator so that, on balance, the system is stable in the absence of diffusion. Mathematically, this extra requirement leads to the inequality

$$\frac{\partial f}{\partial A} < -\frac{\partial g}{\partial I}. \tag{A4}$$

The second constraint we require is that the interactions between the two species be sufficiently strong and have a net stabilising effect in order for the inhibitor to be able to regulate the production of the activator and prevent solution blow-up. Mathematically, this reads

$$0 < -\frac{\partial f}{\partial A}\frac{\partial g}{\partial I} < -\frac{\partial f}{\partial I}\frac{\partial g}{\partial A}. \tag{A5}$$

We see from (A5) that $\frac{\partial f}{\partial I}$ and $\frac{\partial g}{\partial A}$ must have opposing signs, but it is not pre-determined which one is positive and which one is negative. This distinction leads to the two types of Turing reaction-diffusion system. The case $\frac{\partial f}{\partial I} < 0$ and $\frac{\partial g}{\partial A} > 0$ corresponds to Type I (activator-inhibitor) systems, in which the activator promotes the production of the inhibitor and the inhibitor inhibits the production of the activator, and the case $\frac{\partial f}{\partial I} > 0$ and $\frac{\partial g}{\partial A} < 0$ corresponds to Type II (depletion) systems, in which the activator inhibits the production of the inhibitor and the inhibitor promotes the production of the activator (Figure 1C).

The above inequalities, (A3) to (A5), ensure that the system is stable in the absence of diffusion. However, in order for diffusion to drive symmetry breaking, we require that the inhibitor diffuses sufficiently faster than the activator, so that the system exhibits short-range activation and long-range inhibition. Mathematically, this reads

$$D_I > \lambda D_A, \tag{A6}$$

for some constant factor,

$$\lambda = \frac{\left(\sqrt{\det \mathbf{J}} + \sqrt{-J_{12}J_{21}}\right)^2}{J_{11}^2}, \tag{A7}$$

which we have written in terms of the Jacobian matrix of the system for brevity,

$$\mathbf{J} = \begin{pmatrix} J_{11} & J_{12} \\ J_{21} & J_{22} \end{pmatrix} = \begin{pmatrix} \frac{\partial f}{\partial A} & \frac{\partial f}{\partial I} \\ \frac{\partial g}{\partial A} & \frac{\partial g}{\partial I} \end{pmatrix}. \tag{A8}$$

Additionally, since we are investigating a system on a finite ring, we require that the patterns which form as the result of symmetry breaking are sufficiently small so that they fit on our ring. This may be guaranteed by ensuring that diffusion around the centriole surface is sufficiently slow, since if diffusion occurs too quickly then the two species spread evenly around the centriole and there is possibility for patterns to form as the solution becomes uniform. The general criterion for this condition to be met is,

$$\frac{J_{11}D_I + J_{22}D_A + \sqrt{(J_{11}D_I + J_{22}D_A)^2 - 4D_A D_I \det \mathbf{J}}}{2D_A D_I} > 1, \tag{A9}$$

however, a sufficient, yet significantly simpler, condition reads

$$2D_A D_I < \frac{\partial f}{\partial A} D_I + \frac{\partial g}{\partial I} D_A. \tag{A10}$$

Equations (A3), (A4), (A5), and (A9) (or (A10)) describe the conditions required in order to ensure symmetry breaking.

**Appendix II: The activator-conversion model (Model 1)**

The general form of the activator-conversion model, in which the exponents are not specified *a priori*, is given by

$$\frac{\partial A}{\partial t} = \frac{aA^{\alpha}}{1+A} - bAI^{\beta} - cA + D_A \frac{\partial^2 A}{\partial x^2}, \tag{A11}$$

$$\frac{\partial I}{\partial t} = bAI^{\beta} - dI + D_I \frac{\partial^2 I}{\partial x^2}, \tag{A12}$$

for some positive constants $\alpha$ and $\beta$.

Under the appropriate conditions, this system constitutes an activator-inhibitor system as defined in Appendix I, with $A$ acting as the activator, and $I$ acting as the inhibitor. At any non-zero steady state, we find that

$$\frac{\partial f}{\partial A} = (\alpha - 1)\frac{aA^{\alpha-1} + aA^{\alpha}}{(1+A)^2} - \frac{aA^{\alpha}}{(1+A)^2}, \qquad \frac{\partial g}{\partial I} = (\beta - 1)d, \tag{A13}$$

and it is immediately clear that we require

$$\alpha > 1, \qquad \beta < 1, \tag{A14}$$

in order to satisfy (A3). For simplicity, we prescribe $\alpha = 2$ and $\beta = 1/2$ in the main text, and for the remainder of this Appendix, which leads to Equations (3) and (4),

In the case $c = 0$, the two homogeneous steady states of (3) and (4) are given explicitly by

$$(A, I) = (0,0), \qquad (A, I) = \left(\frac{ad - b^2}{b^2}, \left(\frac{ad - b^2}{bd}\right)^2\right). \tag{A15}$$

For the zero steady-state, we find that $\frac{\partial f}{\partial A}, \frac{\partial g}{\partial I} < 0$. Hence, this state does not satisfy (A3) and therefore is unable to break symmetry. For the non-zero steady state, the Jacobian is given by

$$\begin{pmatrix} J_{11} & J_{12} \\ J_{21} & J_{22} \end{pmatrix} = \begin{pmatrix} \frac{(ad-b^2)b^2}{ad^2} & -\frac{1}{2}d \\ \frac{ad-b^2}{d} & -\frac{1}{2}d \end{pmatrix}. \quad \text{(A16)}$$

By substituting this into (A3), (A4), (A5) and (A9), we find that the symmetry breaking conditions are satisfied if

$$0 < ad - b^2 < \frac{ad^3}{b^2}, \quad \text{(A17)}$$

$$D_I > \frac{ad^3\left(\sqrt{ad-b^2} + \sqrt{ad}\right)^2}{2(ad-b^2)b^4} D_A, \qquad D_A D_I < \frac{2(ad-b^2)b^2 D_I - ad^3 D_A}{4ad^2}. \quad \text{(A18)}$$

**Appendix III: <u>The inhibitor-conversion model (Model 2)</u>**

To demonstrate how the reaction-scheme proposed by Leda et al. may be presented within a reaction-diffusion framework, we analyse a modified version of their system of ODE's. Following the notation presented in their manuscript, we focus our attention on the exchange reactions occurring inside the compartments, i.e. the equations for $[PS_i]$, $[P^*S_i]$, $[PS^*{}_i]$, and $[P^*S^*{}_i]$, and we drop the $i$ subscripts for clarity. For the time-being, we will ignore the unbinding/binding terms $R_{10,11,12,14,15}$ since we will reintroduce the concept of compartmental exchange as a diffusion process later. Since they prescribe that $k_1 = k_2 = k_3 = k_4 = k_{-1} = k_{-2} = k_{-3} = k_{-4}$ and $k_5 = k_6 = k_7 = k_8$ in their numerical simulations, we make this simplification upfront in order to simplify the algebra. We label these two reaction rates $\delta$ and $b$, respectively, and note that, in the original model, $\delta \ll b$ when measured over the length-scale of a centriole. However, for the time being, we shall retain the $\delta$ terms. On the other hand, they prescribe that $k_{26}, k_{27} \ll \delta$, and we shall neglect these terms upfront. With these simplifications, the equations read

$$\frac{d[PS]}{dt} = -2\delta[PS] + \delta[P^*S] + \delta[PS^*] - 2b[PS][P^*S^*], \tag{A19}$$

$$\frac{d[P^*S]}{dt} = \delta[PS] - 2\delta[P^*S] + \delta[P^*S^*] - b[P^*S][P^*S^*] + b[PS][P^*S^*], \tag{A20}$$

$$\frac{d[PS^*]}{dt} = \delta[PS] - 2\delta[PS^*] + \delta[P^*S^*] - b[PS^*][P^*S^*] + b[PS][P^*S^*], \tag{A21}$$

$$\frac{d[P^*S^*]}{dt} = \delta[P^*S] + \delta[PS^*] - 2\delta[P^*S^*] + b[P^*S][P^*S^*] + b[PS^*][P^*S^*]. \tag{A22}$$

Summing the species containing phosphorylated STIL complexes and the species containing unphosphorylated STIL complexes yields a system of two equations for $A := [PS^*] + [P^*S^*]$ and $I := [PS] + [P^*S]$,

$$\frac{dA}{dt} = \delta I - \delta A + bI[P^*S^*], \tag{A23}$$

$$\frac{dI}{dt} = \delta A - \delta I - bI[P^*S^*]. \tag{A24}$$

In its current form, the model is underdetermined due to the remaining $[P^*S^*]$ terms. Since the function of these terms is to act as a positive feedback between the phosphorylated STIL complex, $[P^*S^*]$, and the rate at which the unphosphorylated species are converted, we capture this behaviour by replacing $[P^*S^*]$ with a function which depends only on $A$. For simplicity, we prescribe a power law relationship, $[P^*S^*] \approx A^\alpha$, so that the system reads

$$\frac{dA}{dt} = \delta I - \delta A + bIA^\alpha, \tag{A25}$$

$$\frac{dI}{dt} = \delta A - \delta I - bIA^\alpha. \tag{A26}$$

To finalise the model, we must somehow reintroduce a mechanism for transport between receptors along the centriole surface. In the Leda et al., there are a finite number of compartments, and complexes are exchanged between compartments via a shared cytoplasm. In their proposed system, the unbinding rates of $[PS]$ and $[P^*S]$ are orders of magnitude larger than those of $[PS^*]$ and $[P^*S^*]$, since STIL retention at the centriole is postulated to increase with phosphorylation. This results in the species with unphosphorylated STIL being exchanged between compartments via the cytoplasm much more rapidly than their phosphorylated counterparts. In our model, we replicate this effect by assuming that each species diffuses along the surface of the centriole, with the diffusivity of the inhibitor, $D_I$, being greater than the diffusivity of the activator, $D_A$. Since a proportion of the species will be lost to the cytoplasm after unbinding, rather than rebinding and diffusing along the centriole surface, we also introduce additional terms to represent degradation/dissociation. Finally, since the Plk4 and STIL complexes initially bind from the cytoplasm in an unphosphorylated state, we include a constant source term for $I$. Hence, our system reads

$$\frac{\partial A}{\partial t} = \delta I - (\delta + c)A + bIA^{\alpha} + D_A \frac{\partial^2 A}{\partial x^2}, \tag{A27}$$

$$\frac{\partial I}{\partial t} = a + \delta A - (\delta + d)I - bIA^{\alpha} + D_I \frac{\partial^2 I}{\partial x^2}, \tag{A28}$$

for some constants $a$, $c$ and $d$, which correspond to the constant source term and the degradation/dissociation rates of the activator and inhibitor, respectively. Since it is assumed that the activator has a greater retention rate than the inhibitor, we also assume that $c \leq d$.

Equations (A27) and (A28) constitute a reaction-diffusion system whose symmetry-breaking characteristics are determined by the constraints outlined in Appendix I. In general, this must be done computationally. However, in order to gain some insight into the properties of this system, we consider a simplified specific case. Since

$$\frac{\partial f}{\partial A} = -\frac{\delta I}{A} + (\alpha - 1)bIA^{\alpha-1}, \tag{A29}$$

it follows that we require that $\alpha > 1$ in order to satisfy (A3). Therefore, we prescribe $\alpha = 2$. In addition, since (A3) requires that $\delta$ be sufficiently small, $\delta < (\alpha - 1)bA^{\alpha}$, we prescribe $\delta = 0$ to simplify the analysis. This assumption is consistent with our prior note that $\delta \ll b$, and is equivalent to assuming that the only conversion between the two species is due to the non-linear positive feedback phosphorylation term. Since the $\delta$ terms correspond to linear exchange between the two species, this has little effect on the qualitative results.

Our simplified system (which we present in the main paper) therefore reads,

$$\frac{\partial A}{\partial t} = bIA^2 - cA + D_A \frac{\partial^2 A}{\partial x^2}, \tag{A30}$$

$$\frac{\partial I}{\partial t} = a - bIA^2 - dI + D_I \frac{\partial^2 I}{\partial x^2}, \tag{A31}$$

and the diffusion-free steady states are given by

$$A = \begin{cases} 0 \\ \dfrac{ab \pm \sqrt{a^2b^2 - 4bdc^2}}{2bc}, \end{cases} \qquad I = \frac{a}{d} - A. \tag{A32}$$

At the steady state $(A, I) = (0, a/d)$, we find that $\frac{\partial f}{\partial A} = -c < 0$. This contradicts (A3), and therefore this state does not break symmetry.

For the non-zero (with respect to $A$) steady states to exist, we require that

$$ab > 2\sqrt{bdc^2}. \tag{A33}$$

If this condition is satisfied, the Jacobian at these states is given by

$$\begin{pmatrix} J_{11} & J_{12} \\ J_{21} & J_{22} \end{pmatrix} = \begin{pmatrix} c & bA^2 \\ -2c & -d - bA^2 \end{pmatrix}. \tag{A34}$$

Hence, we find that (A4) is automatically satisfied due to the fact that $c \leq d$. On the other hand, we find that (A5) is satisfied if and only if $d < bA^2$. From (A32) and (A33), we find that this is true for the positive root solution, $A = \frac{ab+\sqrt{a^2b^2-4bdc^2}}{2bc}$, and this solution only. Finally, substituting (A32) and (A34) into (A7) and (A10) yields expressions for the remaining constraints,

$$D_I > \frac{\left(\sqrt{Q} + \sqrt{2Q + 4dc^2}\right)^2}{2c^3} D_A, \qquad 2D_A D_I < cD_I - \frac{Q}{2c^2} D_A, \tag{A35}$$

where we have written $Q = a^2b - 4dc^2 + a\sqrt{a^2b^2 - 4bdc^2}$ for clarity. Thus, this model breaks symmetry about the positive-root non-zero steady state, and this state only, provided that the parameters satisfy (A33) and (A35).

**Appendix IV: Coupling diffusion with cytoplasmic concentration**

Since PLK4 degrades in the cytoplasm due to destructive dimerization, it is reasonable to suppose that the "effective" diffusivities of the two species will decrease as the cytoplasmic concentration of PLK4 increases. To model this effect, we define the variable-diffusion functions $D_A^*(a) = a^* D_A / a$ and $D_I^*(a) = a^* D_I / a$, which we substitute into the models in place of $D_A$ and $D_I$, respectively. The normalisation parameter, $a^*$, must be selected appropriately – we prescribe this to be the value of $a$ used in the main simulations (500 for Model 1, 100 for Model 2). With these substitutions, Model 1 reads

$$\frac{\partial A}{\partial t} = \frac{aA^2}{1+A} - bA\sqrt{I} - cA + \frac{500\, D_A}{a} \frac{\partial^2 A}{\partial x^2}, \tag{A36}$$

$$\frac{\partial I}{\partial t} = bA\sqrt{I} - dI + \frac{500\, D_I}{a} \frac{\partial^2 I}{\partial x^2}, \tag{A37}$$

and Model 2 reads

$$\frac{\partial A}{\partial t} = bIA^2 - cA + \frac{100\,D_A}{a}\frac{\partial^2 A}{\partial x^2}, \tag{A38}$$

$$\frac{\partial I}{\partial t} = a - bIA^2 - dI + \frac{100\,D_I}{a}\frac{\partial^2 I}{\partial x^2}. \tag{A39}$$

### Appendix V: Numerical simulations

In the numerical simulations, spatial discretisation is performed using a finite differences scheme with periodic boundary conditions, and time-stepping is performed with the variable-order MATLAB function ode15s. Continuum simulations are performed with 50 grid points, and simulations with $N$ compartments are performed with $N$ grid points. All simulations are deterministic; however, it is straightforward to append a stochastic noise term to the models. Numerical studies with noise reveal that it has negligible impact on the solution and that all qualitative aspects remain consistent with our findings. However, since it complicates both the mathematical analysis and the presentation, so we have chosen to not to include these terms for clarity.
